# Cross-Species Prediction of Histone Modifications in Plants via Deep Learning

**DOI:** 10.1101/2025.05.19.655006

**Authors:** Tongxuan Lv, Quan Han, Yilin Li, Chen Liang, Zhonghao Ruan, Haoyu Chao, Ming Chen, Dijun Chen

## Abstract

The regulation of gene expression in plants is governed by complex interactions between *cis*-regulatory elements and epigenetic modifications such as histone marks. While deep learning models have achieved success in predicting regulatory features from DNA sequence, their cross-species generalizability in plants remains largely unexplored. In this study, we systematically evaluated the ability of deep learning models to predict histone modifications across plant species using a multi-stage framework based on the Sei architecture. We trained species-specific models for Arabidopsis (*A. thaliana*), rice (*O. sativa*), and maize (*Z. mays*), achieving high within-species predictive performance (AUROC > 0.94) and strong agreement between predictions and experimental ChIP-seq profiles. However, cross-species predictions showed reduced performance with increasing phylogenetic distance, highlighting limited model transferability between monocots and dicots. To improve generalization, we built family-level models by training on multiple species across the Poaceae and Brassicaceae families. These models demonstrated robust predictive power in completely unprofiled species—those entirely unused in training set—highlighting the model’s adaptability to novel plant genomes based solely on conserved regulatory syntax. In contrast, cross-family models produced inconsistent results, with reliable performance only in species sharing conserved regulatory features. Lastly, we developed an easy-to-use pipeline that predicts genome-wide chromatin signals directly from DNA sequences. Our findings demonstrate that phylogenetically informed model training significantly improves cross-species epigenomic prediction, offering a scalable computational strategy for functional annotation in non-model and agriculturally important plants.

## Introduction

The precise regulation of gene expression in plants is jointly controlled by complex interactions between DNA sequence elements (such as transcription factor binding sites (TFBS), promoters, enhancers, and inducible elements) and epigenetic modifications (particularly histone modifications and DNA methylation)[1–3]. These elements and modifications are collectively referred to as cis-regulatory elements. Among these, histone modifications — including methylation (e.g., H3K4me3) and acetylation (e.g., H3K27ac) — play a key role in regulating chromatin accessibility and transcriptional activity, thereby significantly influencing plant development, environmental adaptability, and phenotypic plasticity[1,4,5].

In recent years, machine learning strategies — especially deep learning methods — have emerged as the state-of-the-art approach, achieving significant success in predicting various genomic features[6], including cis-regulatory elements[7–10], chromatin states[11,12], and gene expression levels[13,14]. These advancements have significantly enhanced the ability to infer regulatory activity directly from DNA sequences, thereby opening new avenues for understanding genomic function[15,16].

Despite these advances, the field still faces two major challenges. First, there is a lack of systematic evaluation of the cross-species generalization ability of deep learning models[17–20]. Most existing models are trained and validated in a single-species context and have not been evaluated for their transferability to phylogenetically distant plant lineages. Second, the vast majority of deep learning-based regulatory models are developed and evaluated using data from animal systems, humans, or a few model plant species[19,21]. Therefore, their applicability to diverse and agriculturally important plant species remains largely unexplored[22,23]. The complexity of plant gene regulation and evolutionary diversity further exacerbate these challenges[24]. Due to adaptive evolution and the diversification of regulatory mechanisms, regulatory elements and histone modification patterns may exhibit significant differences across plant species[25]. Consequently, deep learning models trained on a single species or closely related species may demonstrate limited generalization ability when predicting genome-wide regulation in distantly related species.

To address the limited cross-species transferability of deep learning models in plant regulatory genomics, we adopted and tailored the Sei deep learning framework — a multi-task, sequence-based architecture originally designed to predict thousands of regulatory features directly from DNA sequences[26]. Previous studies have demonstrated the applicability of the Osei model — a Sei variant — in plant systems, validating its potential for plant-specific regulatory prediction[27]. Leveraging its capacity to integrate diverse regulatory signals, we retrained the model using plant-specific epigenomic datasets to evaluate its predictive performance across multiple species. We hypothesised that training on phylogenetically related or taxonomically diverse species would enhance generalization to untrained lineages. To test this, we developed a four-stage experimental framework comprising: (1) species-specific modeling, (2) cross-species prediction, (3) within-family generalization, and (4) cross-family evaluation. This systematic design enabled a comprehensive assessment of how sequence-based models capture regulatory logic and transfer across evolutionary scales in plants.

## Results

As illustrated in Figure 1A, we implemented this framework using chromatin profiling datasets of histone modifications, collected from multiple plant species in the ChIP-Hub database[28]. For each species, signal tracks corresponding to the same chromatin feature were filtered and unified to identify high-confidence regulatory intervals. A 1,024-bp genomic window centered on each region was extracted and one-hot encoded for model input. We trained the Sei model using a hierarchical stack of convolutional layers followed by multi-task classification to learn multi-scale sequence features associated with chromatin activity. The resulting species-specific models were first evaluated within their respective species and subsequently applied to cross-species settings, including both within-family (e.g., Poaceae, Brassicaceae) and cross-family generalization. The following sections detail the performance outcomes and biological insights derived from each stage.

**Figure 1.**
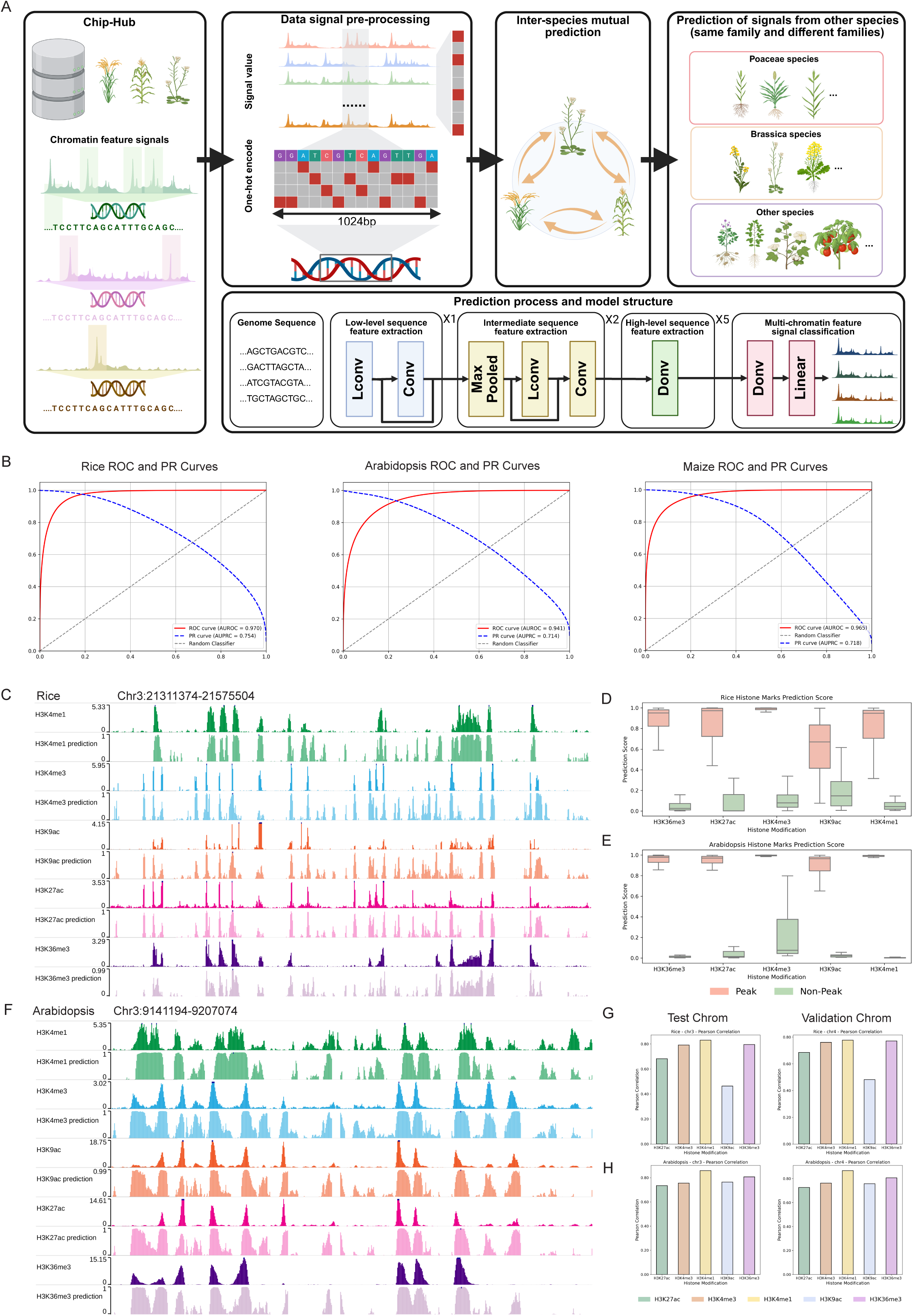
Framework and performance of species-specific chromatin feature prediction in plants. (**A**) Overview of the cross-species chromatin feature prediction framework. (**B**) Receiver operating characteristic (ROC) and precision-recall (PR) curves for species-specific models in Rice, Arabidopsis, and Maize. ROC curves are shown in red; PR curves are shown in blue. (**C**) Predicted and experimental signal tracks for representative histone marks in Rice (Chr3:21311374–21575504). (**D**) The distribution Boxplot of prediction scores for peak and non-peak regions across five histone marks in Rice. (**E**) The distribution Boxplot of prediction scores for peak and non-peak regions across five histone marks in Arabidopsis. (**F**) Predicted and experimental signal tracks for representative histone marks in Arabidopsis (Chr3:9141194–9207074). (**G**) Pearson correlation coefficients between predicted and experimental signals across test and validation chromosomes for each histone mark in Rice. (**H**) Pearson correlation coefficients between predicted and experimental signals across test and validation chromosomes for each histone mark in Arabidopsis.

### Species-Specific Modeling of Chromatin Features in Arabidopsis, Rice, and Maize

We first constructed species-specific models using chromatin feature datasets derived from three representative plant species: rice (*Oryza sativa*), maize (*Zea mays*), and Arabidopsis (*Arabidopsis thaliana*). These species represent major lineages of monocotyledons and dicotyledons and possess rich ChIP-seq datasets, thereby allowing us to evaluate model performance across evolutionarily divergent taxa.

As shown in Figure 1B, the trained models exhibited high accuracy in within-species prediction of chromatin features. Specifically, the rice model achieved an AUROC of 0.970 and an AUPRC of 0.754; the Arabidopsis model achieved an AUROC of 0.941 and an AUPRC of 0.714; and the maize model achieved an AUROC of 0.965 and an AUPRC of 0.718. All performance metrics were calculated on held-out test chromosomes — chromosome 3 for rice and Arabidopsis, and chromosome 6 for maize — ensuring rigorous separation between training and evaluation sets.

To further assess the resolution and biological fidelity of the model predictions, we compared the output signal profiles to ground-truth ChIP-seq tracks from the ChIP-Hub database. As illustrated in Figures 1C, 1F, and 4A, predicted signals in representative genomic intervals—rice (Chr3:21,311,374–21,575,504), Arabidopsis (Chr3:9,141,194–9,207,074), and maize (Chr6:75,059,101–75,148,699)—closely matched experimentally measured chromatin features. To quantitatively evaluate signal specificity, we further conducted peak-versus-non-peak classification by selecting top-ranked peak regions and pairing them with flanking regions of equal length as negative controls. As shown in Figures 1D, 1E, and Supplementary Figure 4B, the models consistently assigned significantly higher scores to peak regions, demonstrating strong discriminatory power across multiple histone modifications, including H3K36me3, H3K27ac, H3K4me3, H3K4me1, and H3K9ac.

To explore the latent regulatory information captured by the model, we performed unsupervised clustering on the predicted chromatin profiles of 1,024-bp genomic windows (Supplementary Figure 1B,2B,3B). To assess the model’s ability to distinguish fine-grained regulatory features beyond histone marks, we also incorporated transcription factor binding site (TFBS) data into this analysis. Using t-SNE for dimensionality reduction, we observed distinct clusters in two-dimensional space, indicating that the model learns biologically meaningful differences in regulatory sequence composition. Enrichment analysis based on ChIP-seq peak annotations further revealed that these clusters corresponded to specific chromatin states: promoter-like clusters were enriched for H3K4me3 and H3K27ac, enhancer-like clusters for H3K4me1 and transcription factor binding, and heterochromatin clusters for CENH3 and H3K9me2 (Supplementary Figure 1A,2A). Based on dominant chromatin features and genomic context, we annotated these clusters into functional categories such as promoters, enhancers, centromeric heterochromatin, and general heterochromatin. In maize, certain clusters showed marked enrichment for individual transcription factors, enabling finer-grained annotation by dominant TF identity (Supplementary Figure 3A).

Lastly, we quantified the Pearson correlation between predicted and experimentally measured signal intensities on both test and validation chromosomes (Figures 1G, 1H, and Supplementary Figure 4C). The majority of histone marks exhibited correlation coefficients exceeding 0.7, indicating that the model reliably captures not only the presence of regulatory features but also their quantitative variation across distinct chromatin marks and genomic partitions. These results underscore the robustness and consistency of model performance across both signal types and genome contexts.

### Cross-Species Prediction of Histone Modifications Using Species-Specific Models

In the second phase of our study, we evaluated the cross-species generalization ability of the species-specific models. Specifically, each model trained on one of the three species (Arabidopsis, rice, or maize) was used to predict histone modification signals in the other two species. This setup allowed us to assess how well regulatory features learned from one species can be transferred to phylogenetically distinct plant genomes. As illustrated in Figure 2A and 2C, we compared the predicted histone modification signals for rice using models trained on maize and Arabidopsis, and conversely, predicted signals for maize using models trained on rice and Arabidopsis. Visual inspection of selected genomic regions revealed substantial differences in signal recovery, with models trained on the more closely related species generally producing signal profiles more consistent with the ground truth.

**Figure 2.**
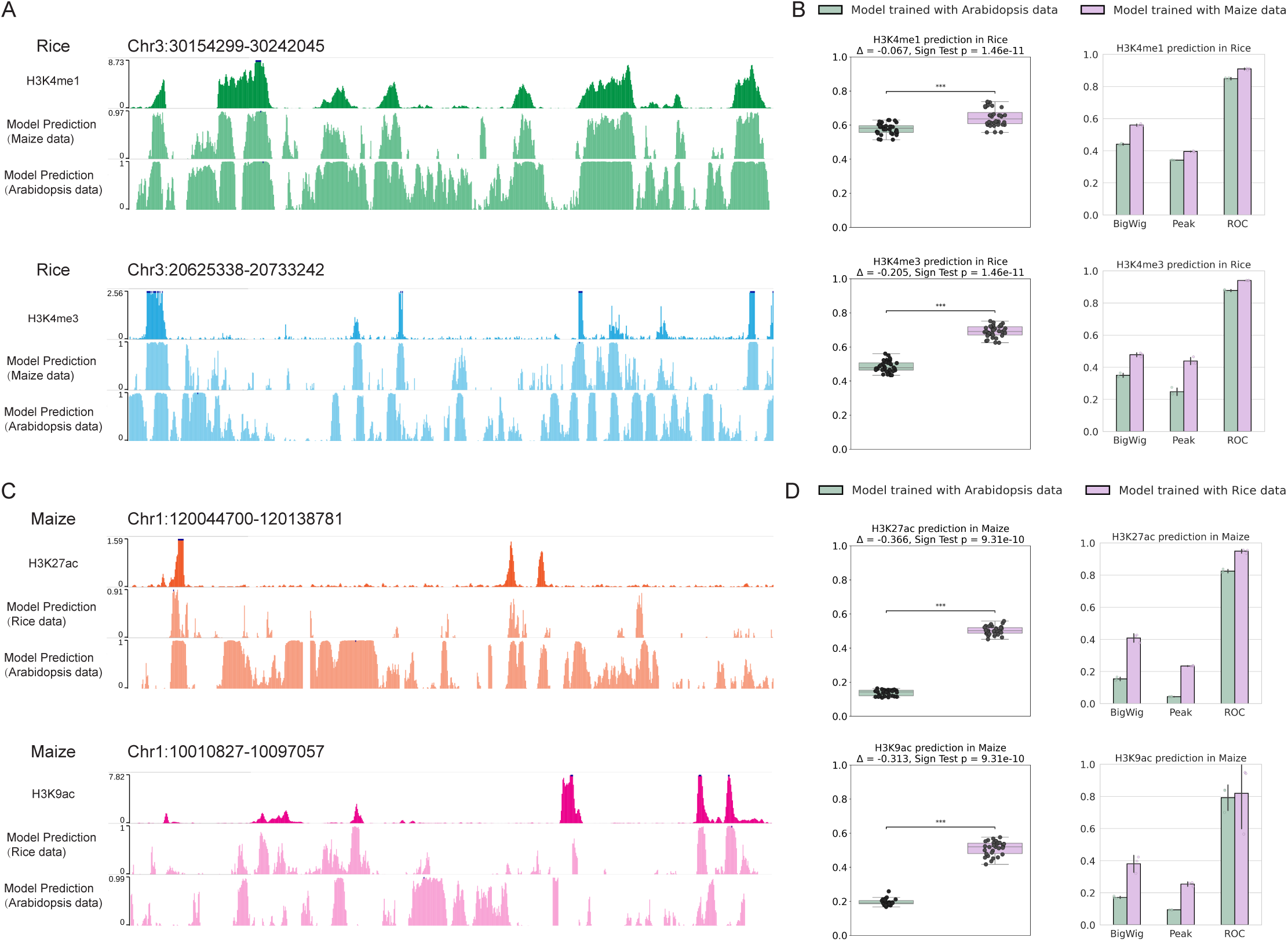
Cross-species prediction performance of chromatin feature models across plant genomes. (**A**) Representative genome browser views showing predicted versus observed signal tracks for histone modifications in Rice using cross-species models. (**B**) Boxplots and barplots showing cross-species prediction scores and evaluation metrics for H3K4me1 and H3K4me3 in Rice using Maize- and Arabidopsis-trained models. (**C**) Representative genome browser views showing predicted versus observed signal tracks for histone modifications in Maize, using cross-species models. (**D**) Boxplots and barplots showing cross-species prediction scores and evaluation metrics for H3K27ac and H3K9ac in Maize using Rice- and Arabidopsis-trained models.

To quantitatively evaluate cross-species prediction performance, we assessed multiple metrics, including Pearson correlation between predicted and observed values across test chromosomes, BigWig-based genome-wide signal correlation, peak-region enrichment, and ROC-AUC scores (Figures 2B, 2D). For rice targets, the model trained on maize consistently produced more accurate signal profiles than the one trained on Arabidopsis, as evidenced by higher correlation coefficients and improved discrimination between peak and non-peak regions. A similar trend was observed when predicting maize signals: the rice-trained model more closely reproduced ChIP-seq patterns than the Arabidopsis-trained model.

As a representative case, we assessed the prediction of H3K27ac histone modification signals in maize using models trained on rice and Arabidopsis. We selected independent genomic regions with high-confidence H3K27ac ChIP-seq signals as ground truth. The rice-trained model achieved a maximum Pearson correlation of 0.52 and a BigWig correlation of 0.42 with the observed signals, while the Arabidopsis-trained model showed markedly lower values of 0.15 and 0.16, respectively. Similarly, we evaluated the prediction of H3K4me3 signals in rice using models trained on maize and Arabidopsis. Across three representative genomic regions, the maize-trained model achieved a maximum Pearson correlation of 0.72 and a BigWig correlation of 0.49, whereas the Arabidopsis-trained model achieved lower correlations of 0.52 and 0.36, respectively.

These results consistently demonstrate the substantial advantage of training on phylogenetically closer species and further emphasize the limited transferability of regulatory models across more distantly related plant taxa.

### Family-Level Cross-Species Prediction with Group-Trained Models

Building on the observation that models trained on phylogenetically closer species exhibit superior predictive performance, we further explored the potential of family-level model generalization. To this end, we constructed a Poaceae family model by jointly training on filtered data from rice and maize. Considering data availability and richness, the Brassicaceae model leveraged the extensive, high-quality epigenomic datasets available for Arabidopsis thaliana, aiming to maximize predictive reliability and consistency while ensuring robust generalization across closely related but less extensively profiled Brassicaceae species. These models were then applied to predict histone modification signals in both family-related and non-family species.

As shown in Figure 3A, the Poaceae-trained model was used to predict chromatin features in other Poaceae family species, including *Setaria italica* (*S. italica*), *Sorghum bicolor* (*S. bicolor*), *Brachypodium distachyon* (*B. dsistachyon*), and *Hordeum vulgare* (*H. vulgare*). Similarly, the Brassicaceae-trained model was evaluated on related species such as *Brassica napus* (*B. napus*), *Arabidopsis alpina* (*A. alpina*), *Brassica rapa* (*B. rapa*), and *Arabidopsis lyrata* (*A. lyrata*). Figure 3B presents the comparison of predicted signal values with true ChIP-seq signals (such as H3K4me1 and H3K4me3) within fixed chromosomal regions across these species. The visualization demonstrates high consistency between the predicted and true signals, particularly in specific chromosomal regions, confirming the model’s robustness in capturing chromatin feature patterns.

**Figure 3.**
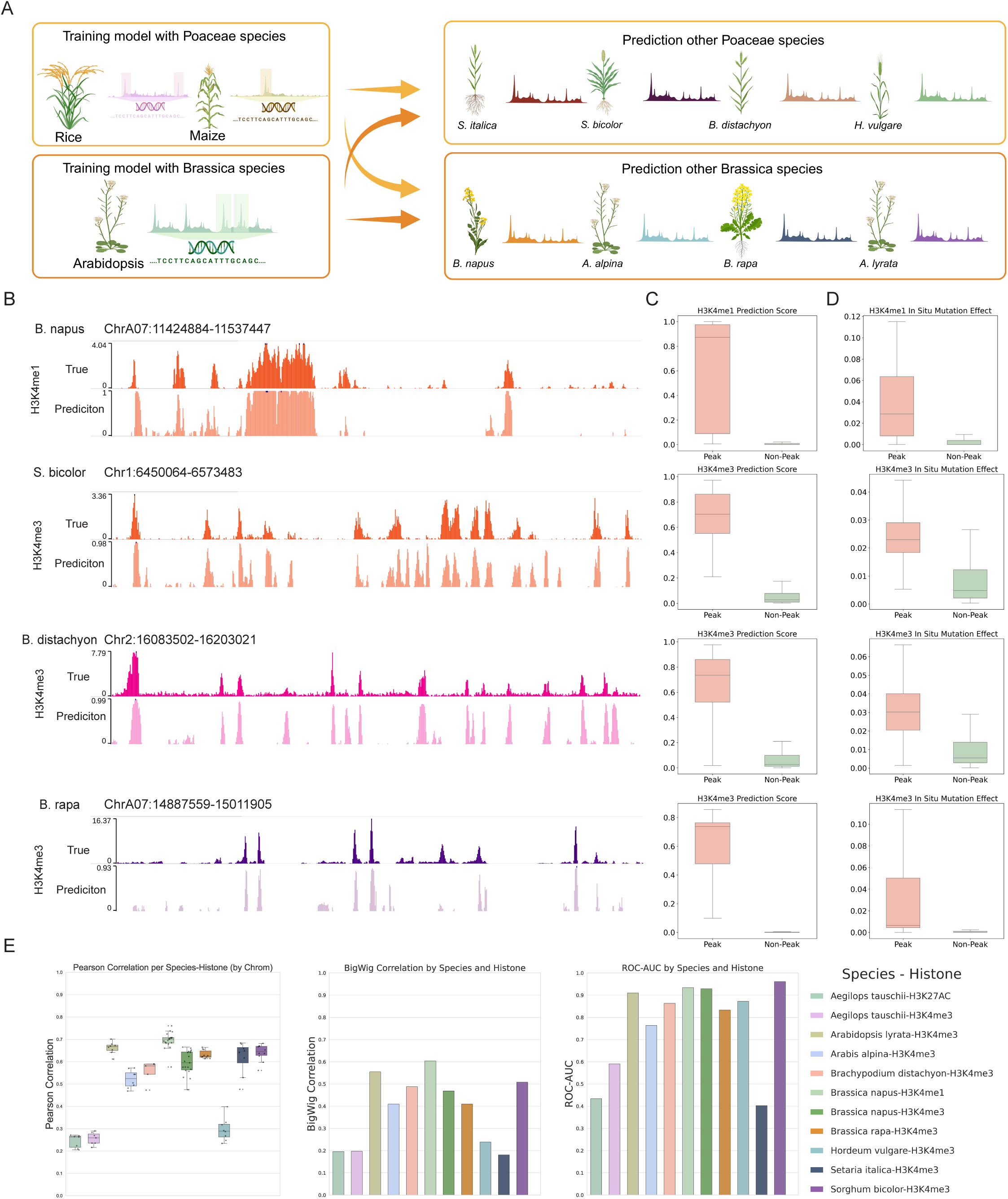
Family-level models improve cross-species prediction of chromatin features. (**A**) Schematic illustration of family-level training and prediction: a Poaceae model trained on Rice and Maize was used to predict signals in other Poaceae species; a Brassicaceae model trained on Arabidopsis was used to predict signals in other Brassicaceae species. (**B**) Comparison of predicted and observed histone modification signals across representative genomic intervals in multiple Poaceae and Brassicaceae species. (**C**) Boxplots showing prediction scores for peak and non-peak regions across three histone modifications. (**D**) Boxplots showing ISM-based Δ scores for peak and non-peak regions across three histone modifications. (**E**) Quantitative evaluation of family-level prediction performance using Pearson correlation, BigWig correlation, and ROC-AUC for each species-histone pair.

To further assess the sensitivity of the family-level models to peak and non-peak regions, we quantified the signal values for both types of regions. Figure 3C shows the prediction results for peak and non-peak regions, with the models consistently assigning higher prediction scores to peak regions, reflecting their ability to accurately capture regulatory signals. In Figure 3D, in silico mutagenesis of important sequence motif (ISM) sites revealed that clipping mutations within peak regions had a significantly higher impact on the predicted signal values compared to mutations in non-peak regions. This further highlights the model’s sensitivity to regulatory features in peak regions. Figure 3E quantifies the correlation between predicted and true signal values, including Pearson correlation, BigWig correlation, and ROC-AUC values, showing that family-level models effectively capture chromatin modification signals for species within the same family.

Additionally, we explored the cross-prediction ability of the Poaceae and Brassicaceae models by using them to predict each other’s species signals, as shown in Figure 4. Compared to within-family models, cross-family models led to false positive peak regions and noise in non-peak regions (Figure 4A). Moreover, the performance of within-family models significantly outperformed cross-family models across different chromosomes, as seen in Figures 4B and 4C.

**Figure 4.**
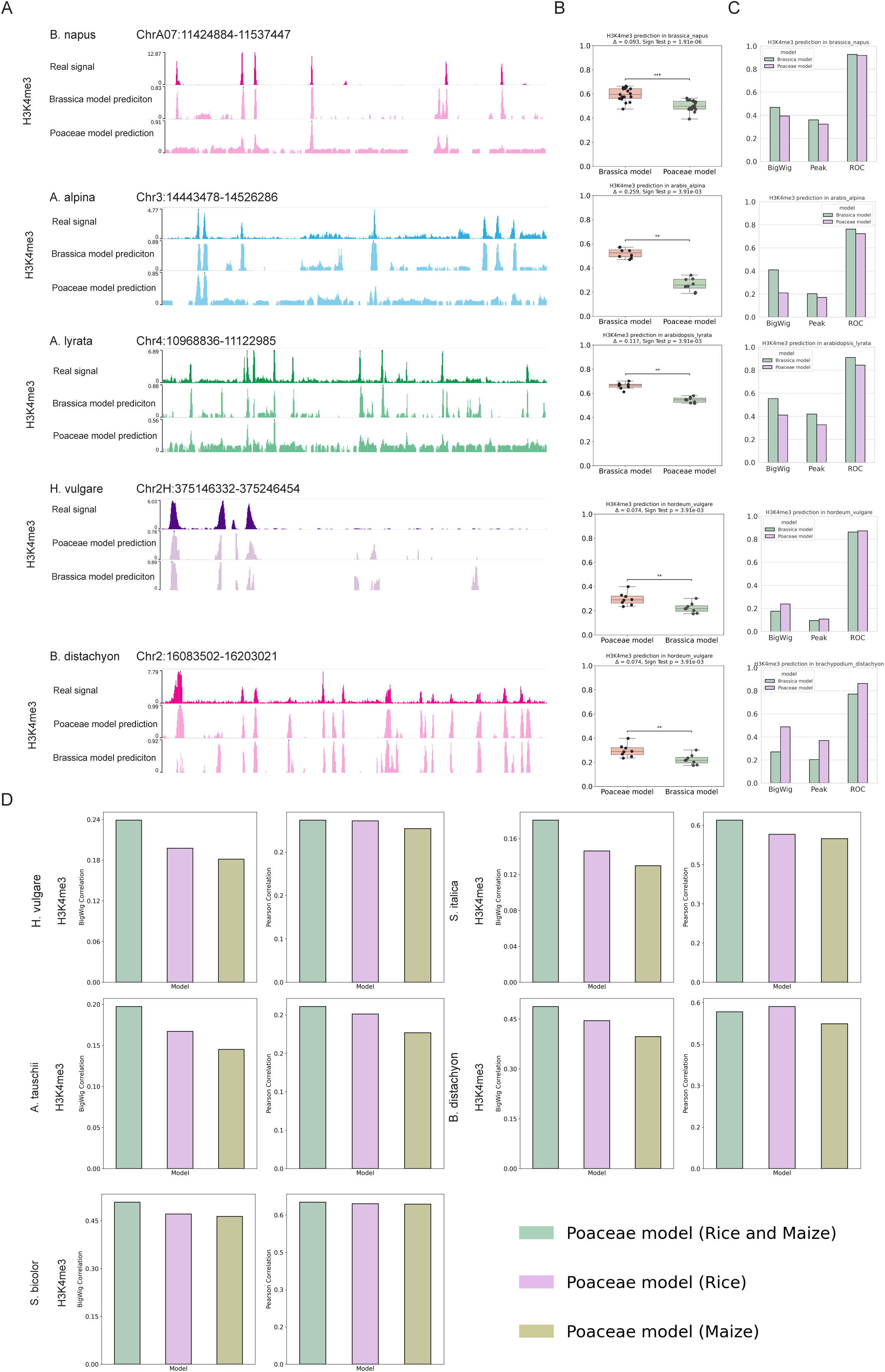
Family-level training improves accuracy and mitigates noise in cross-species prediction. (**A**) Comparison of predicted H3K4me3 signal profiles in five species using family models (Poaceae and Brassicaceae). Each row shows the real ChIP-seq signal, predictions from the Brassicaceae model, and predictions from the Poaceae model across fixed chromosomal intervals. (**B**) Boxplots of prediction scores for H3K4me3 across species, comparing Brassicaceae and Poaceae models. Significance was assessed using the Sign test. (**C**) Bar charts showing BigWig correlation, peak enrichment score, and ROC-AUC for each species, comparing predictions from Brassicaceae and Poaceae models. (**D**) Bar charts comparing model performance across Poaceae species using models trained on Rice only, Maize only, or both combined. Metrics include peak score and average coverage for H3K4me3.

Finally, to evaluate the impact of including multiple species from the same family, we compared the performance of models trained on rice + maize (Poaceae model) with those trained on data from only rice or maize. As shown in Figure 4D, the inclusion of both species significantly improved the model’s performance across various Poaceae species, such as *S. italica*, *S. bicolor*, *B. distachyon*, and *H. vulgare*.

In summary, our results demonstrate that family-level models, trained on phylogenetically related species, significantly enhance the accuracy and generalizability of cross-species chromatin feature prediction, particularly for histone modifications.

### Assessment of Cross-Family Models for Chromatin Feature Prediction Across Diverse Plant Families

To assess whether cross-family models could further enhance predictive performance, we designed the experiment shown in Figure 5A, where we trained a model using filtered signal data from rice, maize, and Arabidopsis to predict chromatin feature signals in both Poaceae and Brassicaceae species, as well as species from other plant families. This multi-family model aimed to determine if incorporating species from multiple families would improve the model’s cross-species prediction capability.

**Figure 5.**
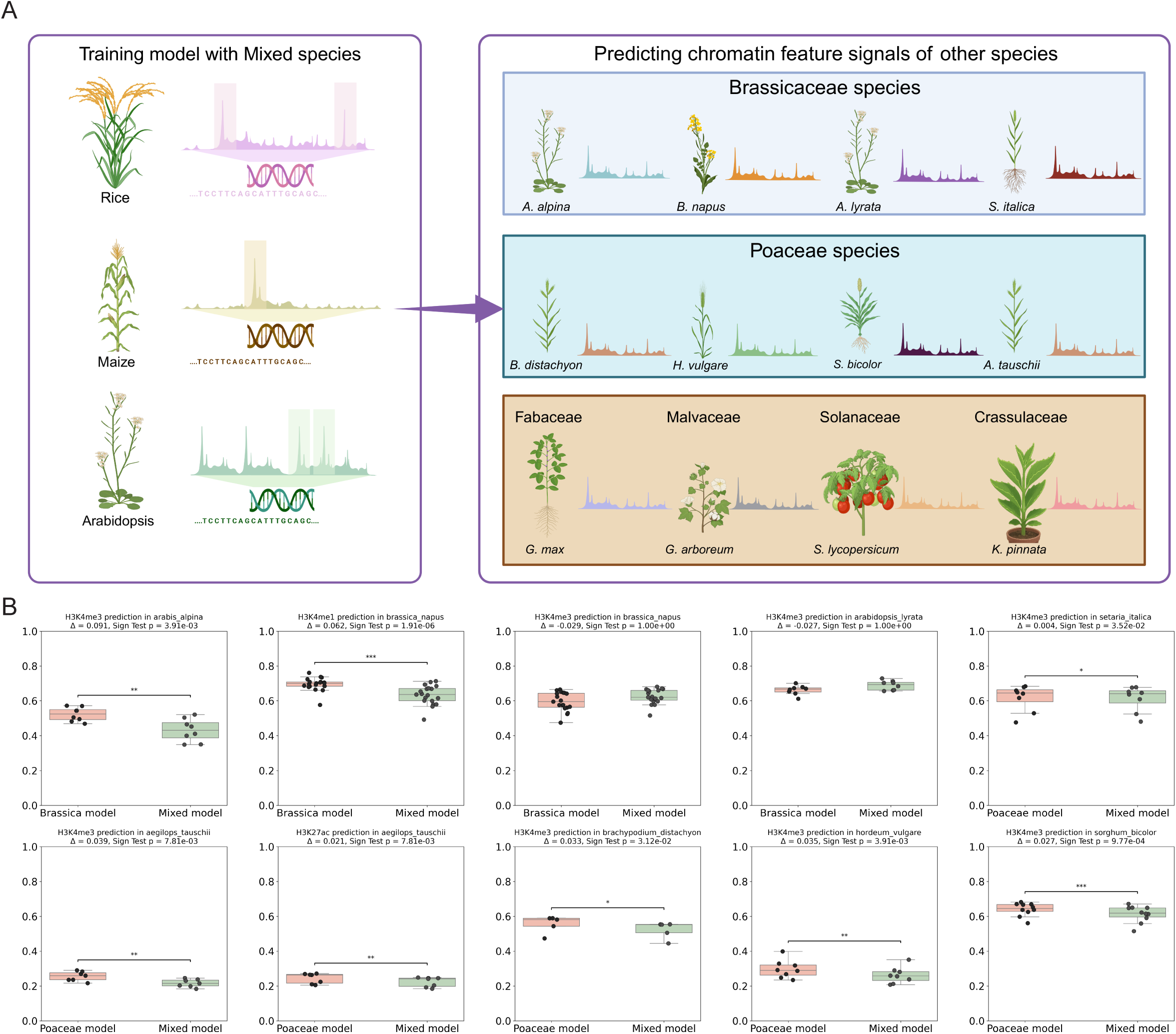
Evaluating cross-family models for chromatin feature prediction across diverse plant lineages. (**A**) Schematic of the cross-family prediction experiment. A multi-species model was trained on Rice, Maize, and Arabidopsis, and used to predict chromatin features in species from Brassicaceae (e.g., *B. napus*, *A. alpina*), Poaceae (e.g., *B. distachyon*, *S. bicolor*), and other families (e.g., *G. max*, *S. lycopersicum*). (**B**) Boxplots comparing family-specific models and the mixed cross-family model across multiple species. Each panel shows prediction scores for a histone mark (mainly H3K4me3 or H3K27ac) in a target species, evaluated using the Sign test. Predictions from the mixed model are shown alongside those from Poaceae or Brassicaceae-specific models.

The cross-family model was trained using a combined dataset of chromatin signals from rice, maize, and Arabidopsis, and was then applied to predict histone modification signals in species from both the Poaceae and Brassicaceae families, including *S. italica*, *S. bicolor*, *B. distachyon*, *H. vulgare*, *B. napus*, *A. alpina*, and *A. lyrata*.

We first evaluated the performance of the cross-family model relative to family-specific models. As shown in Figure 5B, the cross-family model did not consistently outperform the family-specific models. Notably, it showed improved performance in predicting H3K4me3 signals in *B. napus* and *A. lyrata*, but not in other histone marks or species. In contrast, within the Poaceae family, the cross-family model exhibited no performance advantage over models trained exclusively on related species. These results suggest that while cross-family training may offer limited improvements in select cases, especially for specific marks or species, family-specific models generally provide more reliable predictions across most plant taxa evaluated.

We then evaluated the performance of the cross-family model on species from other plant families, including *Glycine max* (*G. max*, Fabaceae), *Gossypium arboreum* (*G. arboreum*, Malvaceae), *Solanum lycopersicum* (*S. lycopersicum*, Solanaceae), and *Kalanchoe pinnata* (*K. pinnata*, Crassulaceae), as shown in Figure 6A and 6B. For histone modification signals such as H3K4me3, H3K4me1, H3K9ac, and H3K27ac, the predicted signals from the cross-family model showed strong correlation with the true signal regions in these species. These results indicate that the cross-family model retains a certain degree of predictive ability for chromatin features even in species from distantly related families. Although the predictive performance of the cross-family model is weaker compared to family-specific models, it still demonstrates a certain ability to accurately capture and localize key signal regions.

**Figure 6.**
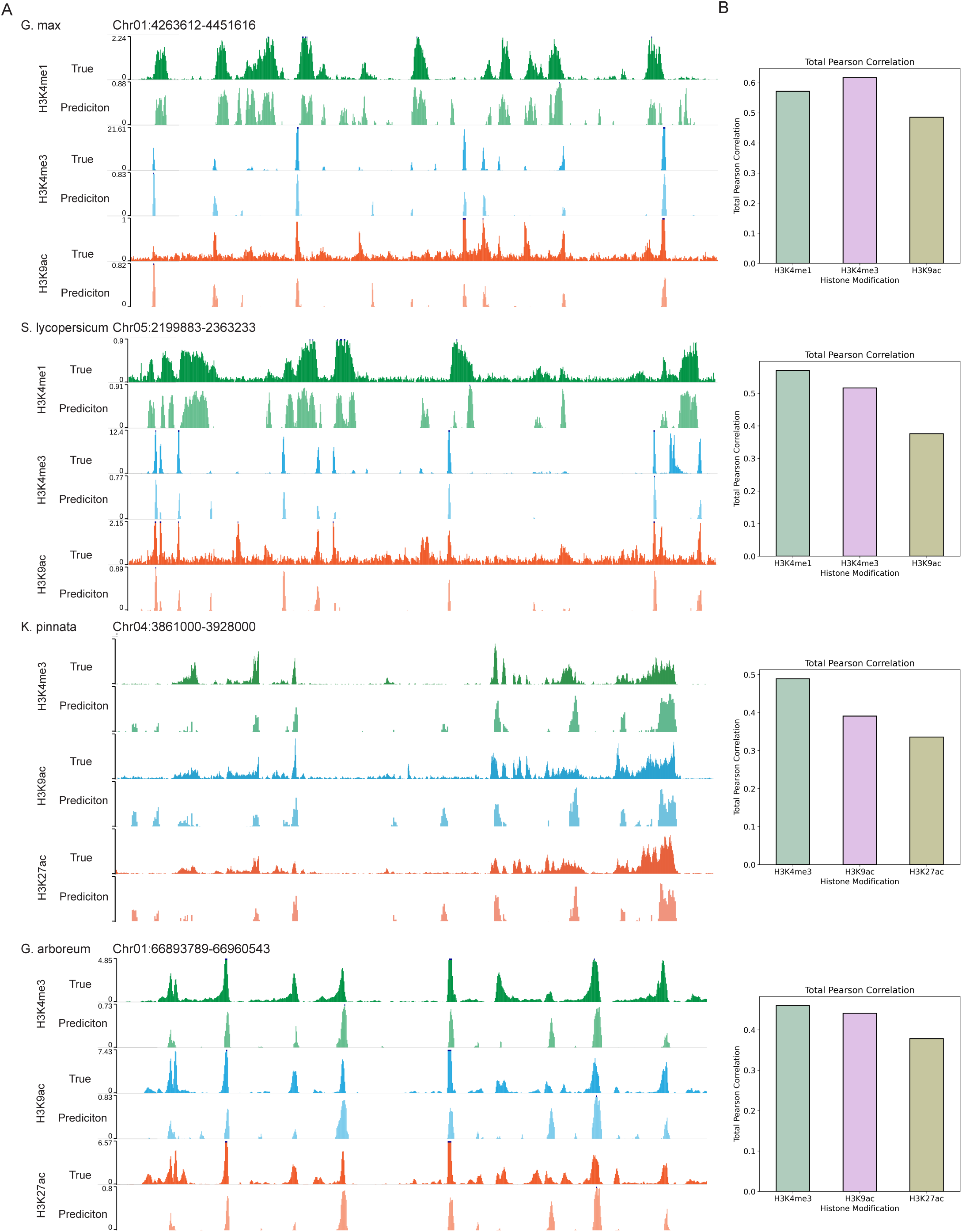
Prediction of chromatin features in non-training lineages using the cross-family model. (**A**) Visualization of predicted versus true ChIP-seq signal tracks in non-training species using the cross-family model. (**B**) Bar plots showing total Pearson correlation between predicted and observed signals for each histone modification.

### Model Usability and Efficient Data Output Generation

To improve model usability, we developed an efficient and user-friendly workflow. As illustrated in Figure 7, users can simply provide the genome sequence file as input, and the model processes it to generate histone modification signal outputs in both BedGraph and BigWig formats. This streamlined, one-click pipeline facilitates the rapid generation of chromatin signal tracks from raw sequence data, allowing for easy integration into downstream analyses. The intuitive interface and accessible output formats significantly reduce the computational burden, enabling researchers to focus on biological interpretation. Importantly, this approach is particularly valuable for species in which chromatin features are difficult or costly to measure experimentally, providing a practical solution for generating epigenomic signal files for rare or under-studied plant species.

**Figure 7.**
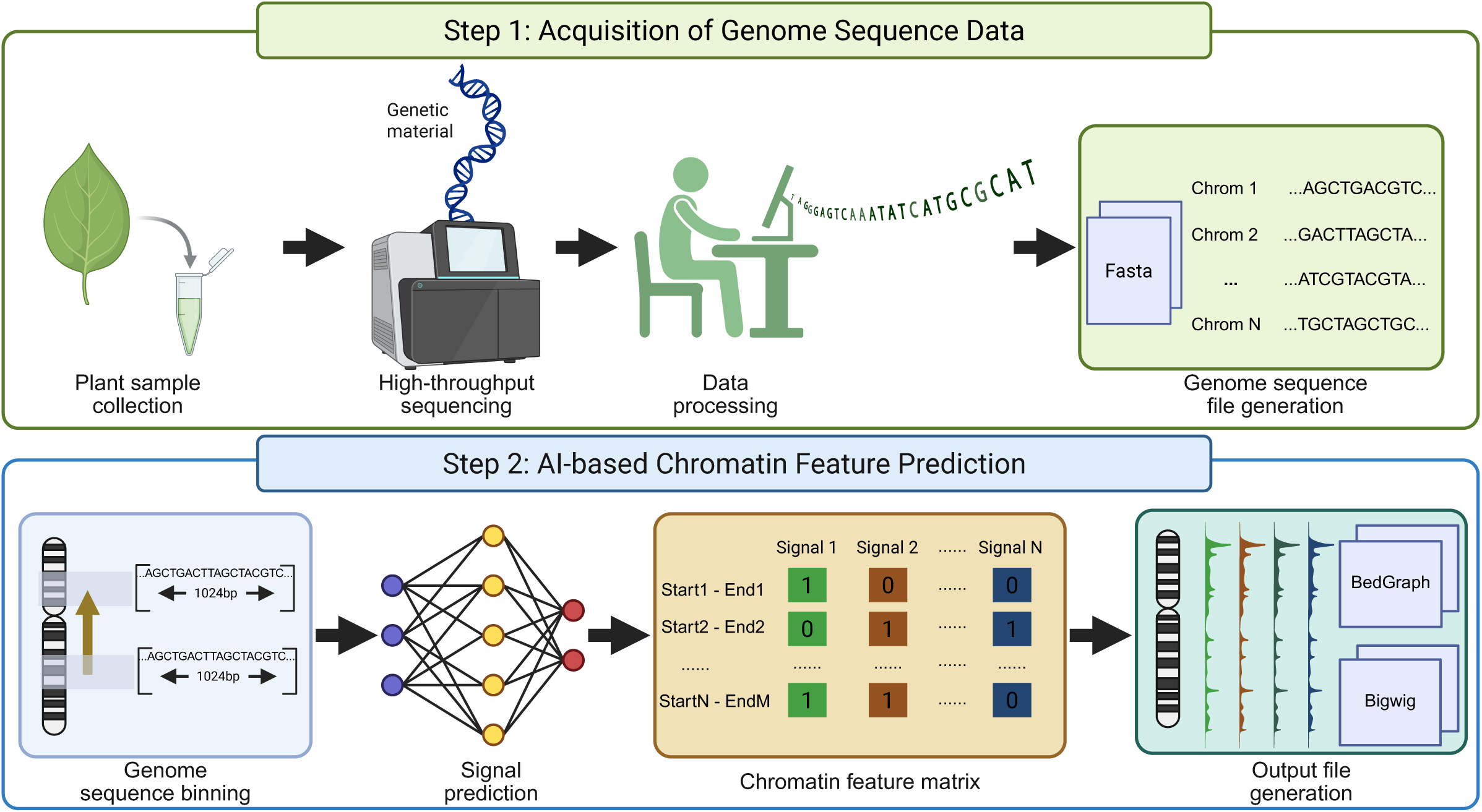
User-friendly pipeline for chromatin signal prediction from genomic sequences. In Step 1, plant samples are collected and sequenced using next-generation sequencing (NGS) to obtain genomic data, which is processed into a standardized FASTA file containing chromosome-level sequences. In Step 2, the genome is binned into fixed-length intervals, and the AI-based prediction model is applied to infer the presence and intensity of histone modification signals.

## Discussion

In this study, we developed and evaluated a deep learning framework for predicting histone modifications across diverse plant species using the Sei architecture—a multi-scale convolutional network originally designed for regulatory sequence modeling. Species-specific models trained on Arabidopsis, rice, and maize achieved strong within-species performance (AUROC > 0.94, AUPRC > 0.71), with predicted signal profiles closely matching ChIP-seq tracks and showing strong enrichment in peak regions. These results demonstrate the model’s ability to decode cis-regulatory logic from raw DNA sequences, consistent with prior findings in human and animal systems[10,19,22,29].

However, model generalization declined with increasing phylogenetic distance. Models trained on monocots (e.g., maize) performed poorly when applied to dicots (e.g., Arabidopsis), and vice versa, highlighting the evolutionary divergence in regulatory syntax between lineages. This phylogeny-dependent loss of accuracy mirrors observations from cross-species chromatin accessibility studies[30], underscoring the challenge of transferring regulatory models across major taxonomic boundaries.

To mitigate this limitation, we trained family-level models using data from multiple species within Poaceae and Brassicaceae. These models significantly improved prediction accuracy in related species, showing higher correlation with experimental data, better peak resolution, and increased robustness. In *silico* mutagenesis further validated the functional importance of sequence features learned by the model, as perturbations within peak regions caused notable drops in predicted signal strength.

In contrast, cross-family models trained on mixed-lineage datasets (e.g., Poaceae + Brassicaceae) yielded inconsistent results. While some conservation was observed — for instance, H3K4me3 predictions in *B. napus* and *A. lyrate* — overall performance lagged behind family-specific models. Interestingly, this improved performance may reflect the relatively high conservation of promoter-associated chromatin features within the Brassicaceae family. Prior comparative studies have shown that regulatory elements marked by H3K4me3 tend to be more conserved among closely related dicots, particularly in the promoters of orthologous genes[31,32]. Such conservation may allow the model to generalize better to these species despite phylogenetic divergence. These findings suggest that although diverse training data may enable broader generalization, phylogenetically informed training remains essential for achieving high-fidelity regulatory predictions in plant genomes with distinct chromatin architectures.

Finally, we developed a streamlined user pipeline that allows users to directly obtain genome browser–compatible visualization files (BigWig and BedGraph) for various predicted chromatin features from raw FASTA sequences. This tool is particularly valuable for species with limited experimental epigenomic data, providing an efficient way to generate interpretable chromatin feature tracks.

Notably, our study does not incorporate additional regulatory layers such as chromatin accessibility (e.g., ATAC-seq)[33,34], transcriptomic output (e.g., RNA-seq), or three-dimensional chromatin architecture (e.g., Hi-C)[35]. Furthermore, the reliance on ChIP-Hub datasets—though high in quality—restricts training to a limited set of histone marks and species with available ChIP-seq data, potentially biasing generalization performance[25].

To improve upon these limitations, future work should explore (i) transformer-based architectures capable of modeling long-range dependencies and integrating prior biological knowledge[21,36,37], (ii) domain adaptation techniques that explicitly account for species differences, and (iii) multimodal models incorporating transcriptomic and chromatin accessibility data. Beyond architectural innovations, expanding the taxonomic breadth and depth of training datasets—especially for under-represented and economically important crops—will be critical to building more inclusive and generalizable models.

## Materials and methods

### Dataset

All data used in this study were obtained from the CHIP-HUB platform (https://biobigdata.nju.edu.cn/ChIPHub/). The data used to train the model consists of two parts: genome sequence data (Fasta format) for each species, and chromatin feature information data (containing histone modification location information and transcription factor binding site location information, in bed and Bigwig formats). The sequence data can be viewed in Table S1. Chromatin feature data for the training of rice, maize, and Arabidopsis species models can be found in Table S2. Chromatin feature data for other Poaceae and Brassicaceae species can be found in Table S3. Chromatin feature data for other species can be found in Table S4. The genomic sequence data and chromatin feature information related to *K. pinnata* were independently sequenced and assembled by our laboratory.

### Training Label Processing

We collected histone modification and transcription factor binding site (TFBS) data for rice, maize, and Arabidopsis from the ChIP-Hub platform, which hosts large-scale, curated chromatin profiling datasets across multiple plant species. For each chromatin feature (e.g., H3K4me3, TFBS), we integrated peak regions from multiple independent ChIP-seq experiments corresponding to the same feature. Genomic segments representing the same chromatin mark across experiments were merged using the “bedtools multiinter” tool in a Linux shell environment. Shared regions detected in at least two independent experiments were retained as high-confidence signals to ensure label robustness and reduce experimental noise. The final set of positive labels was defined as the intersection of these replicates.

For cross-species model evaluation, we focused on five histone modifications with relatively large sample sizes and narrow peak profiles in the ChIP-Hub dataset: H3K4me1, H3K4me3, H3K9ac, H3K27ac, and H3K36me3. To further refine signal confidence, we scored genomic regions based on their overlap frequency across experiments. If all retained regions had exactly two overlaps, they were uniformly assigned a score of 1. Otherwise, we first normalized each region’s overlap count by the total overlap count across all regions, then ranked and linearly mapped the normalized values to a predefined confidence score range (e.g., 0.1 to 1). This final score reflects the consistency of a given region across multiple experimental replicates, with higher scores indicating greater reproducibility and confidence.

### Genome Windowing and Label Generation

After obtaining high-confidence chromatin feature intervals, we loaded the corresponding species’ reference genome sequences. To assign training labels from chromatin features across the genome, we first defined a sliding window approach over the reference genome sequence. To scan the genome in a uniform and non-overlapping fashion, we first defined a sliding window approach over the input genome *G*. Each window *w_j_* spans a 1024-bp interval and slides with a step size of 512 bp. The complete set of genomic windows is defined by:

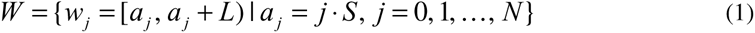

Where *L* = 1024 bp and *S* = 512 bp represent the window size and step size, respectively, and *a_j_* is the starting position of window *w_j_* .

To assign a chromatin activity score to each window, we quantified the total contribution of all overlapping chromatin features derived from BED annotations. Each region (*s_i_*, *e_i_*, *c_i_*) ∈ *R* includes start and end positions *s_i_*, *e_i_* and a confidence score *c_i_* ∈[0,1]. The contribution of region *i* to window *w_j_* is proportional to the fraction of overlap and its confidence score. The aggregated raw score for window *w_j_* is calculated as:

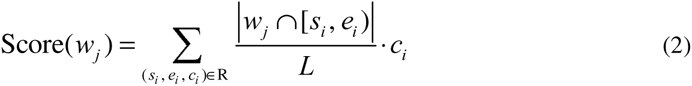

This formula (2) reflects the average per-base contribution of overlapping chromatin signals to the window and effectively integrates signal intensity and genomic span.

For species-specific classification tasks, we generated binary labels for each chromatin feature *k* . A window was labeled as positive if it overlapped any annotated region associated with that feature. Formally, the binary label 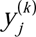 for window *w_j_* is defined as:

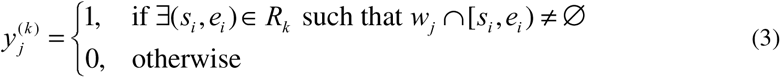

Where *R_k_* denotes the set of genomic intervals associated with histone mark *k*. In cross-species prediction tasks where regression on signal strength is preferred, we normalized the aggregated score for each window to the [0, 1] range. To avoid over-smoothing and to retain interpretability, we rounded each window score to one decimal place and clipped values above 1.0:

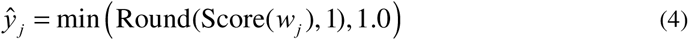

To further suppress noisy or marginal signals that may arise due to weak overlap or low-confidence annotations, we applied a score threshold. Windows with normalized scores below 0.1 were set to zero:

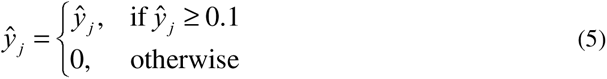

Only windows with non-zero final scores were retained as training candidates in cross-species prediction settings, ensuring that the model focused on informative regions with meaningful regulatory signals.

### Model Architecture and Training Strategy

We adopted a modified version of the Sei model, a multi-task convolutional neural network designed for sequence-based prediction of regulatory features. The model takes as input one-hot encoded 1,024-bp DNA sequences and outputs multi-label predictions corresponding to predefined chromatin features. The architecture includes multiple convolutional blocks for hierarchical feature extraction, followed by parallel dilated convolutions to capture long-range dependencies. A learnable B-spline transformation layer is applied to project the features into a lower-dimensional space, which is then passed to a fully connected classification head. The model was trained using the Adam optimizer with a learning rate of 0.001. For binary classification tasks within a species, we used binary cross-entropy (BCE) loss. For cross-species prediction tasks using continuous signal values, mean squared error (MSE) loss was employed.

### Genome-Wide Signal Prediction and File Generation

To enable genome-wide regulatory signal prediction in non-training species, we implemented a three-step pipeline to generate standardized output files in BedGraph and BigWig formats, facilitating downstream visualization and analysis.

#### Step 1: Genome Sequence Fragmentation

Following model training, we performed genome-wide inference by first generating candidate sequence fragments from the reference genome of the target species. A sliding window approach was applied across each chromosome using a fixed window size (e.g., 1,024 bp) and step size (e.g., 128 bp). Based on the chromosome size file, genomic intervals were systematically extracted and filtered to retain only those containing standard nucleotides (A, T, C, G). The corresponding genomic coordinates were saved in BED format, and the nucleotide sequences were exported in FASTA format to serve as input for downstream prediction.

#### Step 2: Model-Based Signal Inference

Using the pre-trained Sei model, we conducted inference on the unlabeled genomic fragments generated in Step 1. For each input FASTA sequence, the model performs forward propagation to produce multi-label probability scores corresponding to predefined histone modifications. These predicted signal values were saved in NumPy (.npy) format, with each score array aligned to the input window and histone mark.

#### Step 3: BedGraph and BigWig File Assembly

To facilitate visualization and downstream analysis, we assembled genome-wide prediction results into standard BedGraph files. Using the BED coordinates from Step 1 and prediction scores from Step 2, we realigned each window to the central region of the sequence (typically adjusted as start + 448 to end - 576 to match the receptive field center). For each histone mark, the corresponding .npy scores were loaded, filtered, and normalized. Scores below 0.01 were set to zero to suppress weak or background signals. Remaining values were subjected to min-max normalization, mapping signal intensity to a defined range (typically 0.1 to 1.0) while preserving relative magnitude.

The processed scores were then paired with their genomic coordinates to generate per-label BedGraph files. Finally, signal track files were converted to BigWig format using the UCSC utility bedGraphToBigWig, enabling efficient integration with genome browsers and large-scale epigenomic visualization platforms.

### Evaluation of Predicted Signal Accuracy

To quantitatively evaluate the accuracy of predicted histone modification signals, we implemented a multi-metric comparison framework based on both global signal similarity and peak-level agreement between predicted and experimentally measured BigWig files. This evaluation was conducted in three main steps:

#### 1. Genome Binning and Signal Extraction

The reference and predicted BigWig files were processed using pyBigWig to extract average signal values across non-overlapping genomic bins of fixed size (default: 1,024 bp). For each chromosome present in both tracks, signal values were extracted for each bin, and bins containing NaN values were replaced with zero. The extracted values were used to compute global correlation metrics and define peak regions.

#### 2. Global Correlation Metrics

To assess the genome-wide similarity between predicted and experimentally measured histone modification signals, we calculated global correlation metrics using both Pearson methods. The genome was first divided into non-overlapping bins of fixed size (1,024 bp), and the average signal value for each bin was extracted from the predicted and reference BigWig files using the stats() function from the pyBigWig Python package. The resulting signal vectors were then compared using the pearsonr() functions from the scipy.stats module to compute Pearson correlation coefficients. Pearson correlation measures the linear relationship between the predicted and true signal profiles. In addition, for an efficient whole-genome correlation estimate, we employed UCSC’s command-line tool bigWigCorrelate, which directly calculates the Pearson correlation between two BigWig tracks. The output of this tool was recorded as the fast global correlation metric and used as a reference to validate the signal consistency at scale.

#### 3. Peak Region Overlap Analysis

To assess the agreement between predicted and reference histone modification signals in high-signal (peak) regions, we computed the peak overlap ratio based on bin-level peak detection and set intersection. Peak regions were then determined by applying a percentile-based threshold. Specifically, bins whose signal values exceeded the 90th percentile of all bin values in the track were considered peak bins. This process was applied independently to both the predicted signal track and the reference signal track.

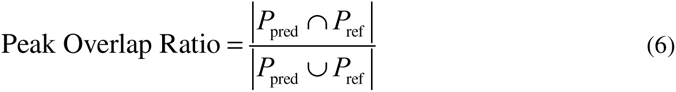

Where *P*_pred_ is the set of peak bins from the predicted signal, *P*_ref_ is the set of peak bins from the reference (ground truth) signal, *P*_pred_ ∩ *P*_ref_ is the number of overlapping peak bins (shared peaks), *P*_pred_ ∪ *P*_ref_ is the total number of unique peak bins across both tracks.

#### 4. ROC-AUC Assessment

To further evaluate the predictive power, we treated the predicted signal values as continuous scores and generated binary labels from the normalized reference signal. Specifically, reference signal values were log-transformed using log1p, followed by min-max normalization. A threshold of 0.5 on the normalized values was used to define binary labels. The ROC-AUC (receiver operating characteristic area under the curve) was computed using sklearn.metrics.roc_auc_score to measure how well the predicted signal distinguished high and low signal regions.

### In Silico Mutagenesis (ISM) and Impact Score Calculation

To assess the functional relevance of individual nucleotide positions within regulatory sequences, we performed in silico saturation mutagenesis (ISM) analysis. Each input sequence (length = 1,024 bp) was first encoded using one-hot encoding (A, T, C, G → 4 channels). For each sequence, we generated all possible single-nucleotide mutations at every position, resulting in 3 × 1,024 = 3,072 mutated variants per sequence. This mutation set included both the reference sequence and all its single-base substitutions. All encoded sequences were passed through a pretrained Sei model to obtain predicted chromatin feature scores. The model outputs, for each sequence or mutation variant, a vector of predicted probabilities across all trained histone modification labels.Then the mutation effect (Δ-score) is defined as:

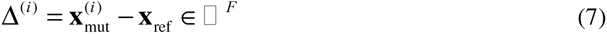

Where *^x^*_ref_ ∈*^F^* is the prediction vector for the reference sequence, 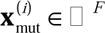 is the prediction vector for the i-th mutation (where *F* is the number of genomic features, e.g., histone marks). For downstream comparison of peak and non-peak regions, we focused on the most impactful mutations by computing the top-5% average absolute Δ-score within each sequence. Specifically, for a target histone mark (indexed by tag), we computed:

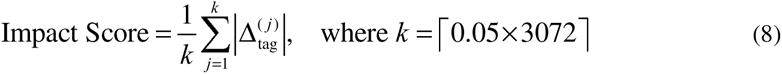

This score reflects the sensitivity of a region’s chromatin feature prediction to small sequence perturbations. Separate impact scores were computed for sequences in peak and non-peak regions.

### Clustering and Functional Annotation of Predicted Chromatin Profiles

Unsupervised clustering was performed on the predicted signal vectors. First, we used t-Distributed Stochastic Neighbor Embedding (t-SNE) for dimensionality reduction to two dimensions. Clustering was then applied to the t-SNE-embedded coordinates to identify distinct groups of regulatory sequences.

To assign functional annotations to these clusters, we performed enrichment analysis utilizing experimentally validated ChIP-seq peak annotations of specific histone modifications and transcription factor (TF) binding sites, including H3K4me3, H3K27ac, H3K4me1, H3K9me2, CENH3, and RNA polymerase (RNAP). Clusters enriched in promoter-associated marks—primarily indicated by RNAP binding sites and H3K4 trimethylation (H3K4me3)—were annotated as promoter regions. Although acetylation marks (such as H3K27ac) are commonly associated with promoters and enhancers, we prioritized RNAP occupancy and H3K4 methylation states for robust promoter annotation due to their specificity and consistent enrichment patterns. Clusters significantly enriched for enhancer-associated marks (H3K4me1 and specific TF binding sites) were annotated as enhancers. Heterochromatic regions were annotated based on characteristic histone modifications: clusters enriched for centromeric-specific CENH3 were labeled as centromeric heterochromatin, clusters enriched for H3K27me3—associated with facultative heterochromatin (regions of reversible chromatin repression)—were annotated accordingly, and clusters enriched for H3K9me2 were categorized as constitutive heterochromatin. For euchromatic regions, the methylation state of H3K4 distinguished enhancer-like regions (monomethylation) from promoter-like regions (trimethylation). In maize, clusters demonstrating strong enrichment of specific transcription factors were further annotated according to the dominant TF identity.

Final functional annotations were systematically assigned based on predominant chromatin features and genomic context, resulting in biologically meaningful categories, including “promoter,” “enhancer,” “centromeric heterochromatin,” “facultative heterochromatin,” and “constitutive heterochromatin.”

## Data And Code Availability

Publicly available datasets used for training and evaluation were obtained from the ChIP-Hub database (https://biobigdata.nju.edu.cn/ChIPHub/). Genomic and chromatin features related to *K. pinnata* (ID: PRJCA040272) can be accessed at: (https://biobigdata.nju.edu.cn/plant2t/upload/36120). The source code for this study is available at GitHub (https://github.com/compbioNJU/SeiPlant). Poaceae, Brassicaceae, and mixed model training parameters and associated configuration files are available at Zenodo: (https://doi.org/10.5281/zenodo.15421964).

## Acknowledgments

We thank the Center for Information Technology and the High Performance Computing Center of Nanjing University for providing computational resources used in this study.

## Author Contributions

DJ.C. designed the research. TX.L. completed the experiments and data preprocessing involved in this paper, completed the first draft of this research paper. Q.H. and ZH.R. assisted in data collection and processing. YL.L. and C.L. assisted in the clustering and annotation of chromatin feature profiles. HY.C. and M.C. assisted in the experimental design of this paper. All the authors reviewed and approved the paper.

