## Supplementary figures and images for "Cross-Species Prediction of Histone Modifications in Plants via Deep Learning"

### Supplemental Figure 1

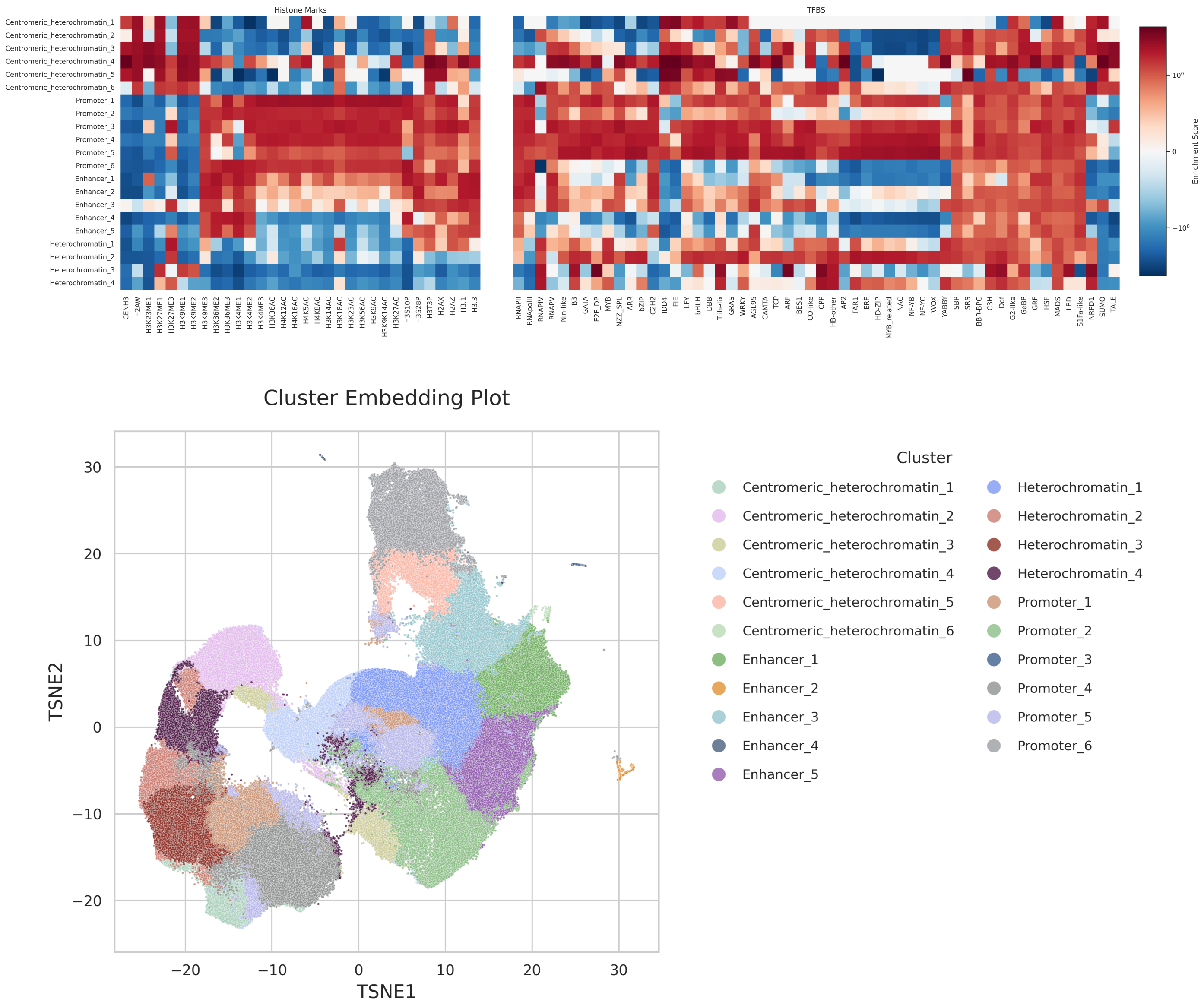

### Supplemental Figure 2

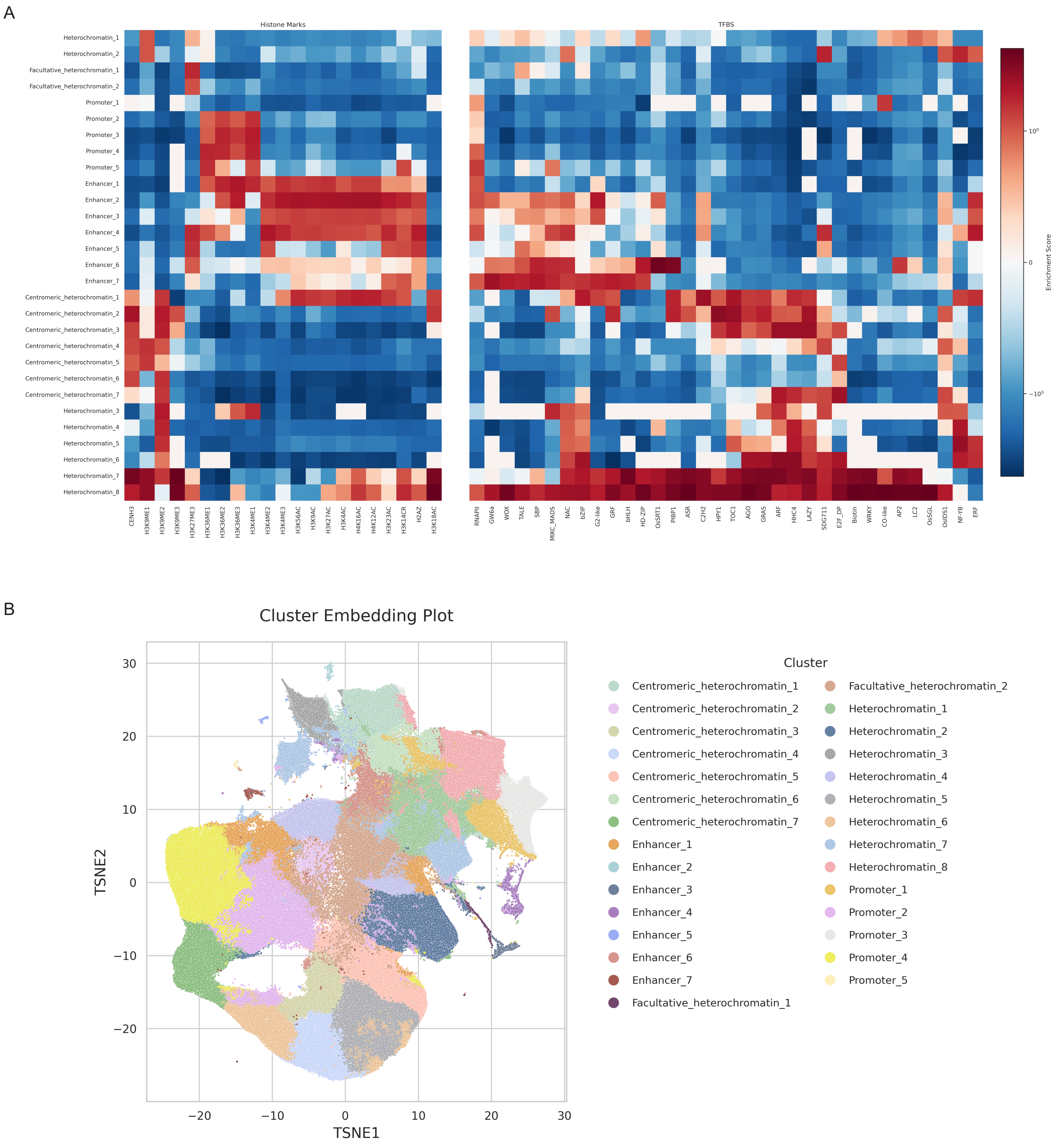

### Supplemental Figure 3

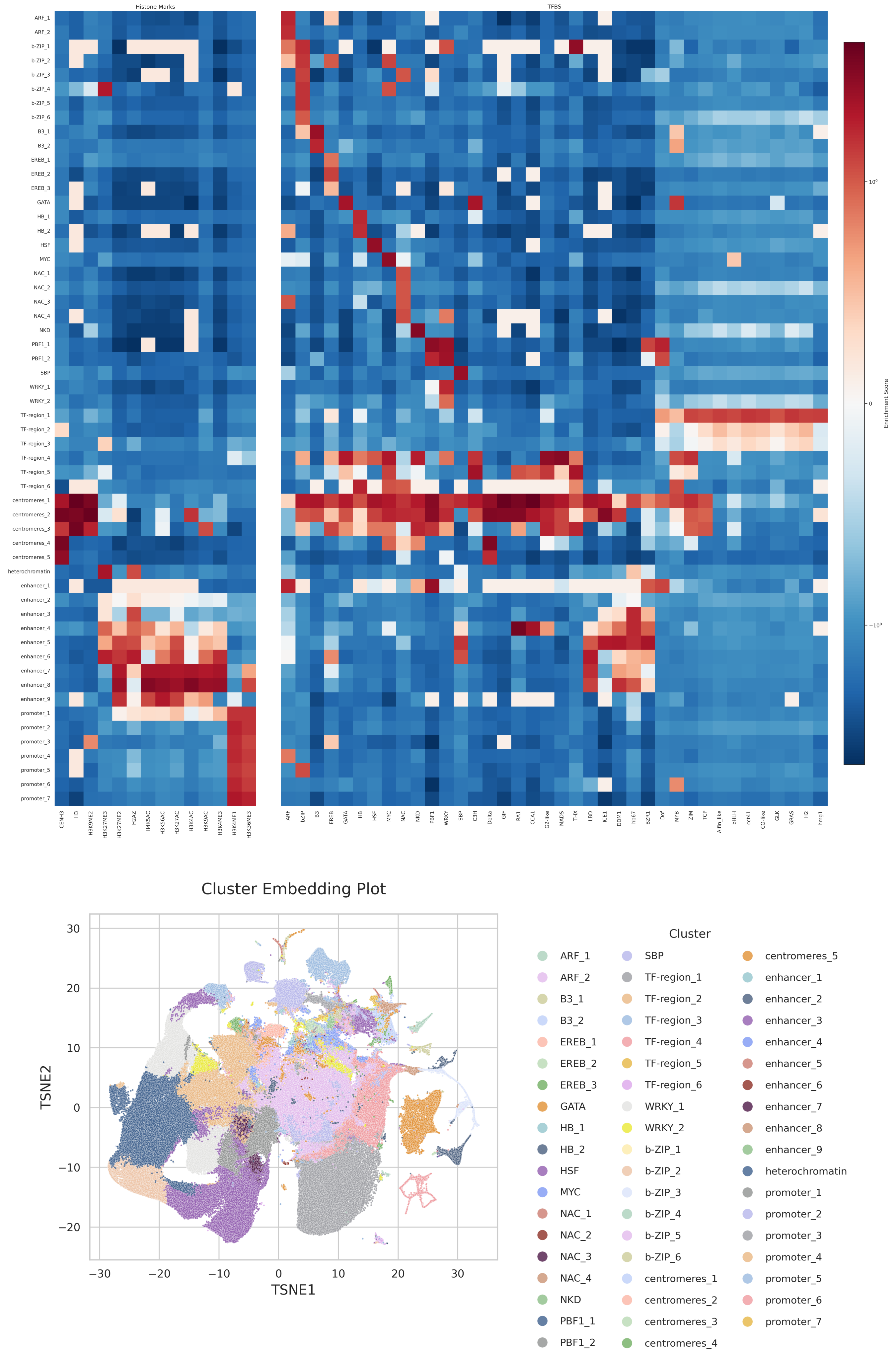

### Supplemental Figure 4

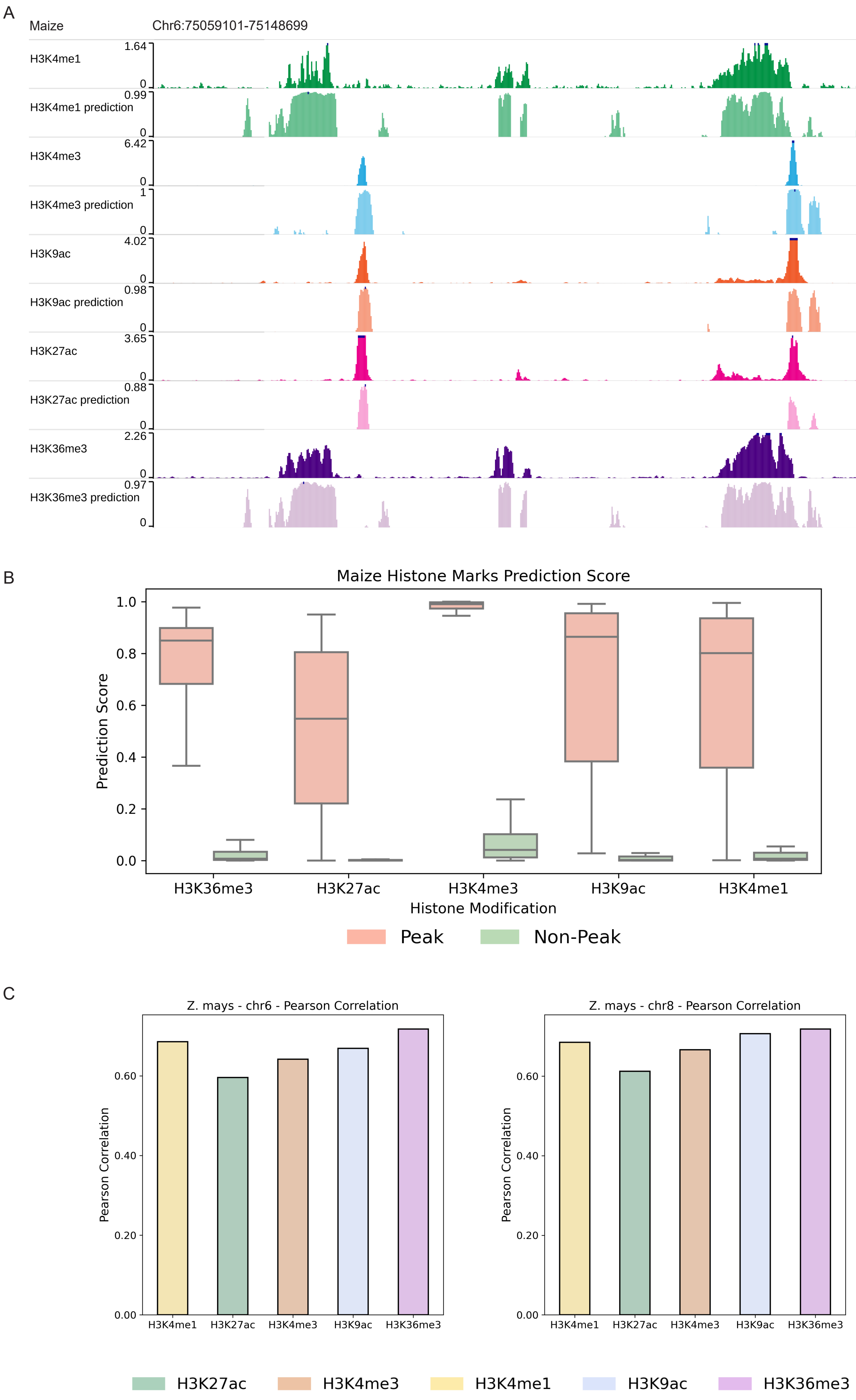
