## Supplemental Table 1-4 for "Cross-Species Prediction of Histone Modifications in Plants via Deep Learning"

**Supplemental Information**

**Supplemental Table 1.** Details of species sequence information.

| Plant species | Common name | Genome release version | Genome size (in Mb) |
| --- | --- | --- | --- |
| *Oryza sativa* | Japonica rice | IRGSP-1.0 | 374.47 |
| *Zea mays* | Maize | AGPv3 | 2066.43 |
| *Arabidopsis thaliana* | Mouse-ear cress | TAIR10 | 119.15 |
| *Arabidopsis lyrata* | Lyre-leaved rock-cress | v1.0 | 200.93 |
| *Arabis alpina* | Alpine rock-cress | v5.0 | 317.91 |
| *Brassica napus* | Rape | v4.1 | 850.29 |
| *Brassica rapa* | Turnip mustard | v1.3 | 297.59 |
| *Setaria italica* | Foxtail millet | v2.0 | 403.32 |
| *Sorghum bicolor* | Sorghum | v3.0.1 | 704.28 |
| *Brachypodium distachyon* | Purple false brome | v3.0 | 271.07 |
| *Hordeum vulgare* | Barley | Hv_IBSC_PGSB_v2 | 4833.79 |
| *Aegilops tauschii* | Tauschs goatgrass | Aet v4.0 | 4078.89 |
| *Glycine max* | Soybean | Wm82.a2.v1 | 949.18 |
| *Gossypium arboreum* | Tree cotton | BGI v2.0 | 1541.29 |
| *Solanum lycopersicum* | Tomato | ITAG2.4 | 823.94 |
| *Solanum tuberosum* | Potato | v4.03 | 773.03 |

**Supplemental Table 2.** Chromatin feature data used for rice, maize, and Arabidopsis

| Species | Histone modification name | Project ID |
| --- | --- | --- |
| *Arabidopsis thaliana* | H3K27AC | SRP251063 |
| *Arabidopsis thaliana* | H3K4ME3 | SRP212174 |
| *Arabidopsis thaliana* | H3K4ME3 | SRP118289 |
| *Arabidopsis thaliana* | H3K4ME3 | SRP075013 |
| *Arabidopsis thaliana* | H3K4ME3 | SRP168223 |
| *Arabidopsis thaliana* | H3K4ME1 | SRP044176 |
| *Arabidopsis thaliana* | H3K4ME1 | SRP336657 |
| *Arabidopsis thaliana* | H3K9AC | SRP152291 |
| *Arabidopsis thaliana* | H3K9AC | SRP227123 |
| *Arabidopsis thaliana* | H3K9AC | SRP170031 |
| *Arabidopsis thaliana* | H3K27ME3 | SRP501366 |
| *Arabidopsis thaliana* | H3K27ME3 | SRP266840 |
| *Arabidopsis thaliana* | H3K36ME3 | SRP193851 |
| *Oryza sativa* | H3K27AC | SRP131213 |
| *Oryza sativa* | H3K27AC | SRP407582 |
| *Oryza sativa* | H3K4ME3 | SRP115109 |
| *Oryza sativa* | H3K4ME1 | SRP238435 |
| *Oryza sativa* | H3K4ME1 | CRA000998 |
| *Oryza sativa* | H3K9AC | CRA009670 |
| *Oryza sativa* | H3K27ME3 | SRP001787 |
| *Oryza sativa* | H3K36ME3 | SRP126351 |
| *Zea mays* | H3K4ME1 | SRP291339 |
| *Zea mays* | H3K4ME1 | SRP195428 |
| *Zea mays* | H3K27AC | SRP158640 |
| *Zea mays* | H3K4ME3 | SRP077759 |
| *Zea mays* | H3K9AC | SRP056839 |
| *Zea mays* | H3K27ME3 | SRP077760 |
| *Zea mays* | H3K36ME3 | SRP162341 |
| *Zea mays* | H3K36ME3 | SRP188687 |

**Supplemental Table 3.** Verified species data sets for Poaceae and Brassicaceae

| Species | Histone modification name | Project ID |
| --- | --- | --- |
| *Arabidopsis lyrata* | H3K4ME3 | SRP036730 |
| *Arabis alpina* | H3K4ME3 | SRP036074 |
| *Brassica napus* | H3K4ME1 | SRP240294 |
| *Brassica napus* | H3K4ME3 | CRA007278 |
| Brassica oleracea | H3K4ME3 | SRP186891 |
| *Brassica rapa* | H3K4ME3 | SRP299560 |
| *Aegilops tauschii* | H3K27AC | CRA005029 |
| *Aegilops tauschii* | H3K4ME3 | SRP094879 |
| *Aegilops tauschii* | H3K4ME1 | CRA009055 |
| *Setaria italica* | H3K4ME3 | SRP158087 |
| *Sorghum bicolor* | H3K4ME3 | SRP110225 |
| *Brachypodium distachyon* | H3K4ME3 | SRP096034 |
| *Hordeum vulgare* | H3K4ME3 | SRP229254 |

**Supplemental Table 4.** Verified species data sets for other family

| Species | Histone modification name | Project ID |
| --- | --- | --- |
| *Glycine max* | H3K4ME1 | CRA012088 |
| *Glycine max* | H3K4ME3 | SRP332511 |
| *Glycine max* | H3K9AC | SRP332511 |
| *Solanum lycopersicum* | H3K27AC | SRP110225 |
| *Solanum lycopersicum* | H3K4ME1 | SRP466644 |
| *Solanum lycopersicum* | H3K4ME3 | CRA015675 |
| *Solanum lycopersicum* | H3K9AC | SRP181007 |
| *Solanum tuberosum* | H3K4ME1 | SRP099193 |
| *Solanum tuberosum* | H3K4ME3 | SRP224921 |
| *Solanum tuberosum* | H3K27AC | SRP394109 |
| *Gossypium arboreum* | H3K4ME3 | SRP114409 |
| *Gossypium arboreum* | H3K9AC | ERP131482 |
| *Gossypium arboreum* | H3K27AC | ERP131482 |
